# Assessing functional connectivity differences and work-related fatigue in surviving COVID-negative patients

**DOI:** 10.1101/2022.02.01.478677

**Authors:** Rakibul Hafiz, Tapan Kumar Gandhi, Sapna Mishra, Alok Prasad, Vidur Mahajan, Benjamin H. Natelson, Xin Di, Bharat B. Biswal

**Author notes:** The authors contributed equally for this project as first authors (Rakibul Hafiz, Tapan Kumar Gandhi).

## Abstract

The Coronavirus Disease 2019 (COVID-19) has affected all aspects of life around the world. Neuroimaging evidence suggests the novel coronavirus can attack the central nervous system (CNS), causing cerebro-vascular abnormalities in the brain. This can lead to focal changes in cerebral blood flow and metabolic oxygen consumption rate in the brain. However, the extent and spatial locations of brain alterations in COVID-19 survivors are largely unknown. In this study, we have assessed brain functional connectivity (FC) using resting-state functional MRI (RS-fMRI) in 38 (25 males) COVID patients two weeks after hospital discharge, when PCR negative and 31 (24 males) healthy subjects. FC was estimated using independent component analysis (ICA) and dual regression. When compared to the healthy group, the COVID group demonstrated significantly enhanced FC in the *basal ganglia* and *precuneus* networks (*family wise error (fwe*) corrected, *p_fwe_ < 0.05*), while, on the other hand, reduced FC in the *language* network (*p_fwe_ < 0.05*). The COVID group also experienced higher fatigue levels during work, compared to the healthy group (*p < 0.001*). Moreover, within the *precuneus* network, we noticed a significant negative correlation between FC and fatigue scores across groups (*Spearman’s ρ = - 0.47, p = 0.001, r^2^ = 0.22*). Interestingly, this relationship was found to be significantly stronger among COVID survivors within the left *parietal lobe*, which is known to be structurally and functionally associated with fatigue in other neurological disorders.

**Significance Statement:** Early neuroimaging studies have mostly focused on structural MRI imaging to report brain abnormalities in acutely ill COVID-19 patients. It is not clear whether functional abnormalities co-exist with structural alterations in patients who have survived the infection and have been discharged from the hospital. A few recent studies have emerged which attempted to address the structural/functional alterations. However, further investigations across different sites are necessary for more conclusive inference. More importantly, fatigue is a highly prevalent symptom among COVID survivors, therefore, the relations of brain imaging abnormalities to fatigue should be investigated. In this study, we try to address these gaps, by collecting imaging data from COVID survivors, now PCR negative, and healthy subjects from a single site – the Indian Institute of Technology (IIT), Delhi, India. Furthermore, this is a continuation of an ongoing study. We have recently shown structural abnormalities and stronger gray matter volume (GMV) correlates of self-reported fatigue in this group of COVID survivors compared to healthy subjects (Hafiz et al., 2022).

## Introduction

The novel coronavirus pandemic has taken more than 4.5 million lives across the globe ((WHO), 2020). Vaccinations and mask mandates have reduced the spread; however, new variants have badly affected densely populated countries, a prime example being, India. Recent evidence shows, Severe Acute Respiratory Syndrome Coronavirus 2 (SARS-CoV-2) can attack the central nervous system (CNS). Clinical MRI from early pandemic reports show cerebrovascular and inflammatory vascular pathologies – acute infarcts, posterior reversible encephalopathy syndrome, microhemorrhages etc. (Gulko et al., 2020; Keller et al., 2020; Nicholson et al., 2020). It is deeply concerning that abnormal fluid-attenuated inversion recovery (FLAIR) uptakes were ubiquitously found in most major brain lobes – frontal, parietal, occipital, temporal, insular and cingulate cortex (Kandemirli et al., 2020), as well as in the sub-cortical systems (Paterson et al., 2020). These clinical cases led to recent studies with more emphasis on group level structural differences (Douaud et al., 2021; Duan et al., 2021; Qin et al., 2021). We have also shown structural alterations in the same group of patients being investigated in the current study (Hafiz et al., 2022). Whether brain pathologies in these survivors also induce functional brain alterations, need to be investigated.

To assess functional brain alterations in COVID survivors, functional magnetic resonance imaging (fMRI) can be implemented. fMRI is sensitive to changes in blood oxygen level dependent (BOLD) signal (Ogawa et al., 1990). Cerebrovascular pathologies in hospitalized COVID patients can modulate blood flow and neural metabolism in the brain. This can lead to abnormal BOLD activity across brain regions, causing alterations in temporal synchronization or functional connectivity (FC) (De Luca et al., 2006; Fox et al., 2005; Fox et al., 2006; Friston, 1994; Friston et al., 1993; Greicius et al., 2003), typically estimated in resting state fMRI (RS-fMRI) among other modalities (Horwitz, 2003). The earliest RS-fMRI study (Biswal et al., 1995) and subsequent studies interpreted FC as information sharing among distinct regions that are temporally synchronized (Cole et al., 2010; De Luca et al., 2006; Fox et al., 2005; Fox et al., 2006; Greicius et al., 2003; Kalcher et al., 2012; Meier et al., 2012). FC can be used to generate robust functional networks (FNs) of the human brain. Independent Component Analysis (ICA) is a popular data driven technique that decomposes the brain voxels into distinct FNs based on the similarity of time courses (McKeown et al., 1998). The ICA-derived large-scale resting-state networks (RSNs) have local and higher level associative hierarchy (Yeo et al., 2011) and replicate highly reproducible activation maps across subjects (Smith et al., 2009).

Distributed brain pathologies in hospitalized survivors can affect multiple FNs. Based on earlier evidence (Matsuda et al., 2004; McCray et al., 2007), it is likely the source of neuro-invasion is the olfactory pathways. The current COVID literature also shows abnormalities in the olfactory system (Esposito et al., 2022; Ismail & Gad, 2021; Laurendon et al., 2020; Niesen et al., 2021; Politi et al., 2020). The virus can subsequently stay and cause pathologies by migrating throughout the brain post infection (Daniel et al., 2022). Therefore, it is possible that most FNs are affected and ideally, they should all be investigated. However, to avoid multiple testing complications and based on our recent study (Hafiz et al., 2022) and current literature findings, we focused on FC alterations primarily in five relevant networks – the *basal ganglia, precuneus*, *language* and *bilateral somatosensory* networks.

The *basal ganglia* is a major hub for projections to and from the cortex and has neuronal connections to the olfactory system (Amunts et al., 2005; Soudry et al., 2011). Recently, we have shown higher GMV within the *basal ganglia* among the same group of survivors in this study (Hafiz et al., 2022). *Precuneus* is a major constituent of *default mode network (DMN*), involved in a multitude of functions including visuo-spatial, memory retrieval, self-referential and switching processes during goal-oriented task (Cavanna & Trimble, 2006; Freton et al., 2014). Reduced FC was reported within *DMN* and between *DMN* and *salience (SAL*) networks in unresponsive COVID survivors (Fischer et al., 2021). Investigating the *precuneus* network may be important for fatigue and attention related deficits among survivors. The *somatosensory* network processes several body related stimuli (Favaro et al., 2012; Kang et al., 2018; Lavagnino et al., 2014) which may relate to symptoms experienced by survivors such as loss in appetite, depression, and sleep disorder among others. The *language* network consists of regions from the *inferior frontal* and *middle temporal gyri (IFG, MTG*). Duan and colleagues have shown structural abnormalities specifically within the *IFG* and *MTG* regions in COVID survivors (Duan et al., 2021). The *language* network also comprises *cerebellar* regions where both structural abnormalities (Fadakar et al., 2020; Malayala et al., 2021), as well as, enhanced dynamic functional connectivity with *sensorimotor* network have been reported among survivors (Fu et al., 2021).

Behaviorally, many COVID survivors experience a sequela of symptoms (Logue et al., 2021; Peluso et al., 2021; Tabacof et al., 2020), now commonly called ‘Long COVID’, which point to brain as the responsible organ. Fatigue, lack of attention, anxiety, memory loss, delayed recovery of smell and/or taste, muscle pain and stress are commonly reported symptoms. Most contemporary neuroimaging studies have mainly focused on brain correlates of post-traumatic stress syndromes (Benedetti et al., 2021; Fu et al., 2021). On the other hand, several others have attempted to use FC as a neurobiological indicator of higher stress levels (Liu et al., 2021; Perica et al., 2021), depression (Zhang et al., 2022) and negative affect (Xiao et al., 2021) among healthy subjects before and after the pandemic. Despite fatigue being the most frequently reported symptom, very little is known of its brain related effects among survivors. In our recent investigation (Hafiz et al., 2022), we also observed a stronger positive correlation between GMV and work-related fatigue within the *precuneus, posterior cingulate cortex* and *superior parietal lobule*. We expect to see similar relationship of FC and work-related within the *precuneus network*.

We first apply ICA in RS-fMRI data to estimate FC in healthy controls (HCs) and COVID survivors using group ICA and dual regression (C.F. Beckmann, 2009; Filippini et al., 2009) to test our hypothesis that surviving COVID-negative patients demonstrate altered FC in the *basal ganglia, precuneus, somatosensory* and *language* networks. Using a self-reported fatigue questionnaire on a scale of 0-5, we further hypothesized that the COVID group is more susceptible to fatigue during work. We also apply a multiple linear regression model to test the hypothesis that FC within the *precuneus* network demonstrate stronger correlation with fatigue compared to HCs.

## Materials and Methods

### Participants

This is a continuation of our recent work using the same sample groups where structural brain alterations were reported (Hafiz et al., 2022). 47 COVID patients and 35 HCs were recruited from a single site located at Indian Institute of Technology (IIT), Delhi, India. 9 COVID and 4 HC subjects were removed during quality control and motion assessment, leaving with an effective sample of 38 (25 males) COVID and 31 (24 males) HC. The COVID subjects were scanned two weeks after they were released from the hospital when confirmed to be COVID-negative upon polymerase chain reaction (PCR) retesting. During scanning, all protocols were strictly followed based on the Institutional Review Board (IRB) guidelines at the Indian Institute of Technology (IIT), Delhi, India.

The patients in this study were recruited from a much larger cohort who were admitted and assessed at the Metro Heart and Super-specialty Hospital, New Delhi, India. The patient evaluation and classification was based on illness severity data derived from a database of 2,538 COVID patients from May to December 2020. 24% of this sample did not require O_2_, 40% required O_2_, 22% required Continuous Positive Airway Pressure (CPAP); and 14% were intubated. This 14% of intubated patients were excluded from the recruitment process in the current study.

Of the remaining patients, 333 needed CPAP to raise O_2_ levels; 333 needed nasal O_2_ to raise O_2_ levels; and 334 were admitted but did not need supplemental O_2_. The sample of 47 COVID subjects constituting the COVID group in this study were collected from this cohort (those who agreed to participate in this ongoing study so far). Patients were recruited two weeks after discharge, after becoming PCR negative. Any subject from the healthy group who has experienced fever, cough or flue like symptoms in the two weeks prior to scan, were removed from the study. All healthy subjects also had to undergo a PCR test to assure that they had not been infected in the recent past. We used a questionnaire to record symptoms the survivors have experienced during hospitalization (see Figure S1 in the Supplementary Materials) and an additional questionnaire to quantify fatigue levels (see Figure S2 in the Supplementary Materials) which also included similar questions from the first questionnaire to identify if they experienced any persistent or new symptoms. Please note this questionnaire (Figure S2) has been used to assess fatigue in patients with Chronic Fatigue Syndrome (CFS) and Fibromyalgia (Natelson, 2019). To avoid confounding effects from comorbidities, we recruited subjects that were, otherwise, in excellent health conditions prior to hospitalization for COVID-19. For example, 16/47 (34.04%) of our patients were young adults who had no prior record of any comorbidities that could confound the COVID-19 effects. We only had 7/47 (14.89%) subjects with age between 50-54 years (capped at < 55 years as recruitment criteria to avoid aging-related comorbidities), some of whom had reported to have marginal diabetes. The rest (51.07%) in between also did not have any record of comorbidities in the hospital report.

*Table 1* summarizes the clinical information from the 47 patients included in the current study. Of these 47, 36.17% (17/47) patients were reported to be ‘mild’, 8.51% (4/47) to be ‘moderate’ and 36.17% (17/47) to be ‘severe’. Information from the rest of the 19.15% (9/47) was not provided from the hospital because those patients did not give consent to sharing their medical symptoms. Please note, we present percentages as a ratio of affected patients with both the available sample with information (‘% Out of Avail.’ in *Table 1*) and the total sample of patients including those patients who did not give consent to publicly share their clinical data (‘% Out of Total’ in *Table 1*). The percentages of ‘moderate-severe’ patients who were administered medications, e.g., Remdesivir, Dexamethasone, Ceftriaxone, Clexane and other antibiotic regimes are provided in *Table 1*. On average these 47 patients stayed in the hospital for approximately 11 ± 3.30[SD] days.

**Table 1.**
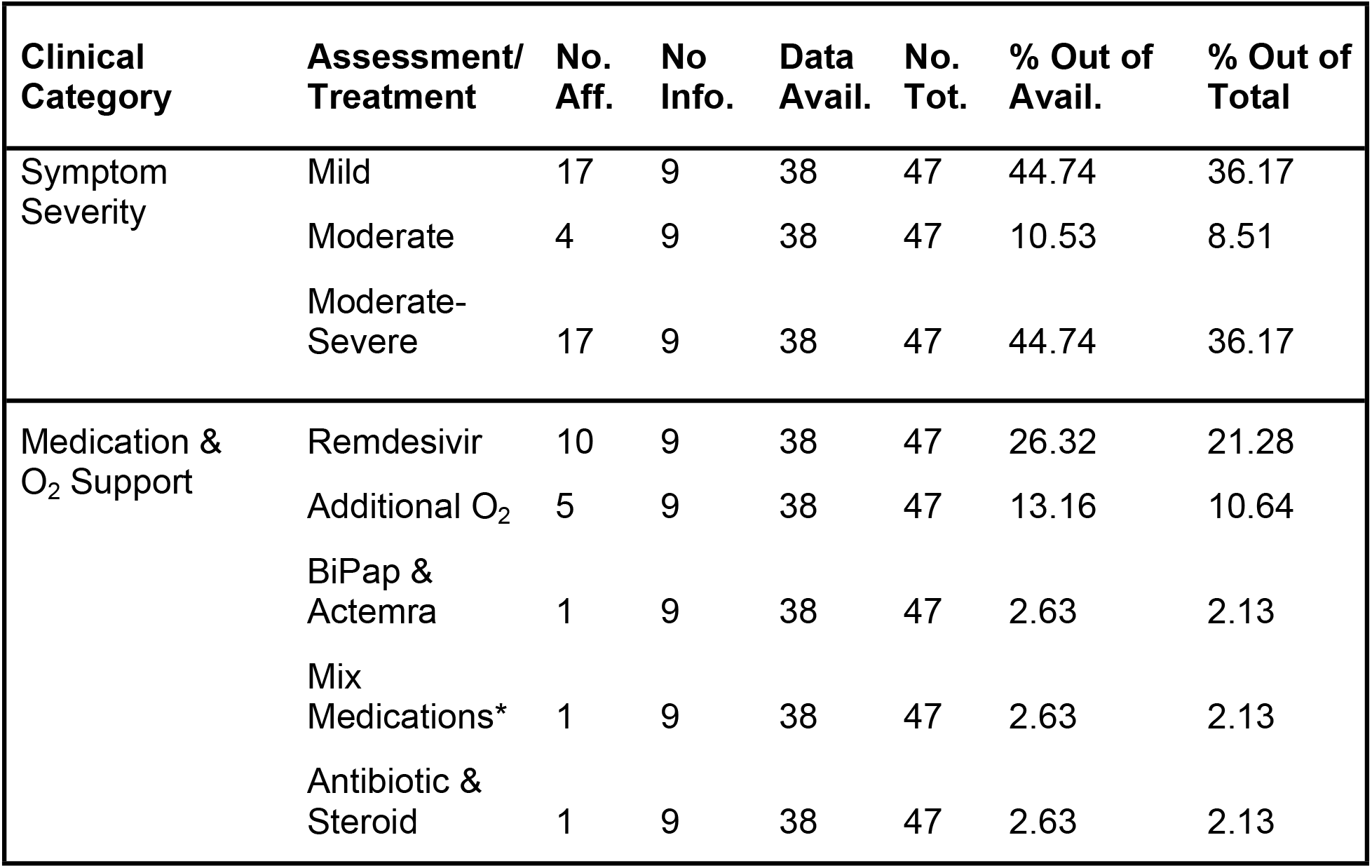
Clinical information detailed by symptom severity and medical treatment of COVID participants. The table shows the number and percentages of participants based on two clinical categories. The first three rows show number and percentage of participants by symptom severity and the last five rows are based on the type of medication administered and the requirement of O_2_ support. Particularly, information from 9 participants could not be obtained as these participants did not agree to share the symptom publicly/anonymously. **Keys:** No. Aff. = number of affected patients, No Info. = no information available because these patients did not give consent to share symptom information, Data Avail. = number of subjects with clinical assessment data available, No. Tot. = total number of patients including those with no information available, % Out of Avail. = proportion of patients affected vs patients with clinical assessment data available (n = 38) in percentages, % Out of Total = proportion of patients affected vs. total number of patients (n = 47) in percentages, O_2_ = oxygen supplied to support breathing, BiPap = bilevel positive air pressure, Mix Medications* = a combination of medications – Dexamethasone, Ceftriaxone and Clexane injections

*Table 2* summarizes the participant demographics based on age, sex and fatigue. The average age in the HC group was 33.50 ± 9.74 [SD] years and that in the COVID group was 34.63 ± 11.54 [SD] years. The number of males was comparatively higher than females in the HC group 23M vs. 7F, however, the same was true for the COVID group 31M vs. 15F. The participants were asked through a questionnaire (see Supplementary Materials, Figure S2), what level of fatigue do they experience during their daily work. Note this ‘work’ is related to their daily profession and not household chores. They were asked to rate their fatigue levels on a scale of 0 to 5, with 0 representing no fatigue and 5 representing the highest fatigue possible. The healthy group underwent the same questionnaire and reported their daily fatigue levels during work. Fatigue scores were successfully obtained from HC (n = 17) and COVID (n = 27) groups. The mean and standard deviations of fatigue scores in the HC group was 0.65 ± 0.79 [SD] and that of the COVID group was 2.93 ± 1.21 [SD].

**Table 2:** Group level statistics on participant demographics among HC and COVID group. The table shows the test results from participant demographics including age, sex and fatigue. There was no significant differences in age and sex between the two groups. However, the COVID group experienced significantly higher fatigue levels compared to HC group. **Keys:** *p = p-value, t = two-sample t-test statistic, χ^2^= Chi-Squared statistic, T = Wilcoxon Rank Sum test score*, M = Male, F = Female.

| Measures | *p* | *stat* | HC, mean (SD) | COVID, mean (SD) |
| --- | --- | --- | --- | --- |
| Age (years) | 0.66 | -0.44 (*t*) | 33.50 (9.74) | 34.63 (11.54) |
| Sex | 0.38 | 0.76 ($\chi^2$) | 23M, 7F | 31M, 15F |
| Fatigue | 2.097e-07 | 36.5 (*T*) | 0.65 (0.79) | 2.93 (1.21) |

### Clinical Assessment

The most frequently reported symptoms from the participants (see Supplementary Materials, Figure S1 for questionnaire used) during hospitalization were - fever, cough, body ache, chills, difficulty breathing, bowel irritation, nausea, loss of sense of smell and loss of consciousness. From the day of discharge till the day of scan, we further asked if the participants were experiencing any persistent or new symptoms. Work-related fatigue (86.84%), muscle pain (18.42), lack of sleep (39.47%), lack of attention (36.84%), headache (36.84%), joint pain (50%), memory loss (34.21%), delayed recovery of sense of smell (44.74%) and/or taste (34.21%), bowel irritation (57.89%) and interestingly, hair loss (81.58%) were commonly reported. Please note, most survivors experienced multiple symptoms simultaneously, hence the ‘%’ represents symptoms that overlap within participants. For example, 36.84% of COVID participants reporting with lack of attention also reported a work-related fatigue score > 2.

### Brain Imaging

#### Anatomical MRI

High-resolution T1-weighted images were acquired on a 3T GE scanner with a 32-channel head coil in 3D imaging mode with a fast BRAVO sequence. The imaging parameters were TI = 450 ms; 244 x 200 matrix; Flip angle = 12 and FOV = 256 mm. The subject was placed in a supine position and the whole brain was scanned in the sagittal configuration where 152 slices were collected, and each slice was 1.00 mm thick. The spatial resolution of all the anatomical scans was 1.0 mm x 1.0 mm x 1.0 mm.

#### Resting-state fMRI

A gradient echo planar imaging (EPI) was used to obtain 200 whole-brain functional volumes. The parameters were: TR = 2000 ms; TE = 30 ms; Flip angle = 90, 38 slices, matrix =64×64; FOV = 240 x 240 mm^2^; acquisition voxel size = 3.75 x 3.75 x 3 mm^3^. The participant was requested to stay as still and motionless as possible with eyes fixed to a cross on an overhead screen. Please note, while initially we kept our scanning time limited to collecting 200 time points due to severe pandemic conditions, as conditions eased, we increased our scanning time to collect 400 time points in this ongoing study.

### Data Pre-Processing

The data preprocessing was performed primarily using Statistical Parametric Mapping 12 (SPM12) toolbox (http://www.fil.ion.ucl.ac.uk/spm/) within a MATLAB environment (The MathWorks, Inc., Natick, MA, USA). However, some steps utilized useful tools from FSL (FMRIB Analysis Group, Oxford, UK) and AFNI (http://afni.nimh.nih.gov/afni) (Cox, 1996) for housekeeping, visual inspection and quality control purposes. At the beginning, first five time points were excluded from each subject to account for magnetic stabilization. The functional images were motion corrected for head movement using a least squared approach and 6 parameters (rigid body) spatial transformation with respect to the mean image of the scan. The subjects with excessive head motion were identified using framewise displacement (FWD) (Power et al., 2012). Additionally, time frames with high FWD crossing a threshold of 0.5 mm (Power et al., 2012) were identified along with the previous and the next two frames and added as regressors (Yan et al., 2016) during temporal regression of nuisance signals. If more than 50% of the time series data were affected due to regression of high motion frames the participant was removed from the analysis. Moreover, any participant with the maximum framewise translation or rotation exceeding 2 mm was removed from further analysis. Anatomical image from each subject was coregistered to the mean functional image obtained from the motion correction step. T1-weighted image from each subject was segmented into gray matter (GM), white matter (WM), and cerebrospinal fluid (CSF) tissue probability maps and an average template including all participants was generated using DARTEL (Ashburner, 2007). The subject specific tissue maps were non-linearly warped to this template and spatially normalized to the MNI space. These affine transformations were applied to the functional images to normalize all volumes to the MNI space and resampled to isotropic voxel size of 3 mm x 3 mm x 3 mm. Time series, from brain compartments with high physiological noise signals such as, CSF and WM was extracted by thresholding the probability maps from the segmentation stage above the 99^th^ percentile, and first 5 principial components were obtained using a COMPCOR based (Behzadi et al., 2007) principal component analysis (PCA) from both tissues. These 10 components along with Friston’s 24-parameter model (6 head motion parameters + 6 previous time point motion parameters + 12 corresponding quadratic parameters) (Friston et al., 1996) and time frames with high FWD (> 0.5 mm) were added as regressors in a multiple linear regression model to remove unwanted signals voxel-wise. The residuals from the regression step were then bandpass filtered between 0.01 to 0.1 Hz and finally, spatial smoothing was performed using a Gaussian kernel of 6 mm full width at half maximum (FWHM).

### Head Motion Assessment

We performed in-scanner head movement assessment using mean Framewise Displacement (FWD) based on the methods depicted in (Power et al., 2012). A two-tailed two-sample student’s t-test revealed no significant differences in mean FWD between the two groups (*t = −1.57, p = 0.12, α = 0.05*).

### ICA and Dual Regression

Group level resting state networks were obtained by applying the ‘gica’ option of the ‘melodic’ module from FSL toolbox (FMRIB Analysis Group, Oxford, UK). All subjects’ 4D functional images after pre-processing were temporally concatenated into a 2D matrix of ‘space’ x ‘time’ as delineated in (C.F. Beckmann, 2009) and 25 spatial maps were obtained. Resting State Networks (RSNs) were identified by matching ICs with the 1000 functional connectome project maps (Biswal et al., 2010) using Dice’s coefficient and spatial correlations obtained from AFNI’s ‘3dMatch’ program (Taylor & Saad, 2013). Further visual inspection was performed to make sure all network regions aligned with the functional network and ROIs depicted in (Altmann et al., 2015; Shirer et al., 2012). Dual regression (C.F. Beckmann, 2009; Filippini et al., 2009) was performed leveraging the standardized group ICA output from the ‘melodic’ step and applying it directly to the ‘fsl-glm’ module in FSL to obtain subject specific RSN maps. The subject specific network maps were standardized to Z-scores before consequently applying them in statistical analysis to infer group level estimates.

### Statistical Analysis

To investigate differences in participant demographics, we performed a two-sample *t-test* on age. Since sex is a categorical variable, we performed a *chi-squared* test to identify any sex related differences between the groups. Since the fatigue scores deviated from normality (*Shapiro-Wilk, p < 0.05*), we performed a non-parametric *Wilcoxon-Ranksum test* on the fatigue scores to identify group level differences in fatigue scores.

To investigate FC differences between COVID and HC groups, we performed an unpaired two sample t-test between standardized subject-specific RSN maps from the two groups. To account for confounding effects that may explain some of the variance in the data, age, sex and a regressor to account for two different scanning lengths were also added as covariates of no interest. Cluster-based thresholding was applied at a height threshold of *p_unc_ < 0.01*, with *family wise error (FWE*) correction at *p_FWE_ < 0.05* for multiple comparisons. The cluster extent threshold (*k_E_*) obtained from this step was used to threshold and generate corrected statistical maps for the contrasts with significant effects.

We further wanted to evaluate if the PRN demonstrates correlation with self-reported fatigue among the COVID individuals. We incorporated a multiple linear regression approach where the FC at each voxel was the response variable (Y), and the self-reported fatigue score was the explanatory variable (X). We also added age and sex as covariates of no interest. Significant clusters were obtained in the same manner as described earlier at the end of the previous paragraph for group level differences in FC. For visual representation of the significant relationship between the two variables, the average FC within the significant cluster was obtained from each subject. These average FC values were then linearly regressed against the fatigue scores and visualized within a scatter plot and a line of best fit with 95% confidence interval. Age and sex were regressed out during the linear regression step. The correlation analysis and the graphical plotting was done using ‘inhouse’ scripts prepared in RStudio (RStudio, 2021).

## Results

We will present results on participant demographics first and then group level voxel-wise results will be reported. There was no significant differences in age between the two groups (*p > 0.05*). A *chi-squared* test on sex revealed no significant effects were observed between the two groups (*p > 0.05*). The Wilcoxon-Ranksum test revealed significantly higher fatigue levels in the COVID group compared to the HC group (*T = 1093, p = 2.86e-07*).

We identified twenty-two large-scale resting state networks (RSNs) (see Figure 1) from the group ICA analysis. Group level statistical analysis was run for five networks of interest using standardized subject specific RSN maps obtained from the dual regression step. Significant differences in FC was observed between the COVID and HC groups in particularly three out of the five networks – the *basal ganglia (BGN), precuneus (PRN*) and *language (LANG*) networks. Figure 2 shows all significant clusters from obtained from the *t*-test and the corresponding group level networks where these alterations occur.

**Figure 1.**
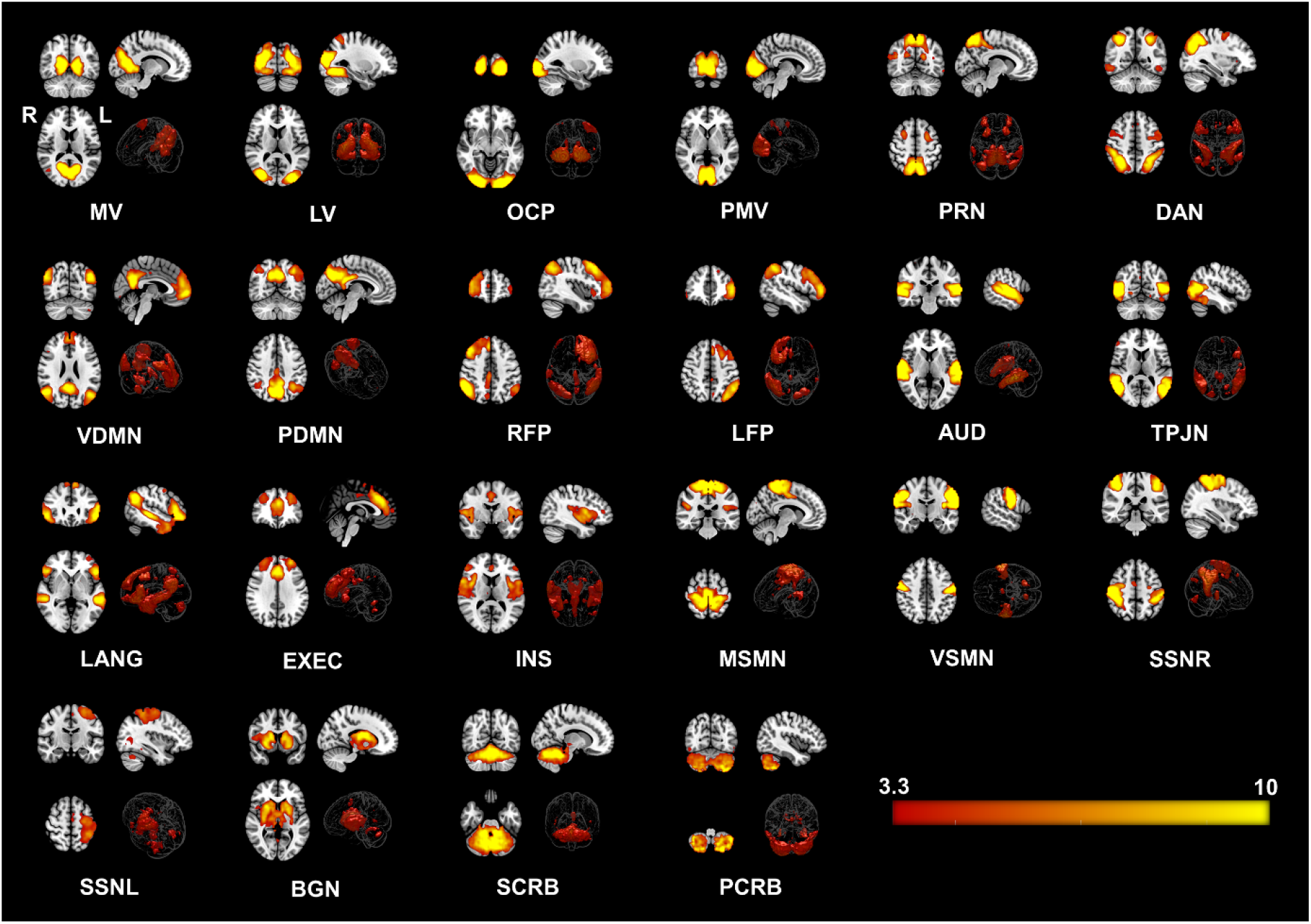
Twenty-two Resting State Networks (RSNs) identified from group ICA using ‘melodic’. Abbreviated names of each network are shown at the bottom of each image. Three orthogonal slices are shown for each network along with a volume rendered image to show depth and three-dimensional view of the RSNs. Statistical estimates (Z-scores) are embedded into a colorbar at the bottom-right. **Keys:** MV = Medial Visual, LV = Lateral Visual, OCP =Occipital Pole, PMV = Primary Visual Network, PRN = Precuneus Network, DAN = Dorsal Attention, VDMN = Ventral Default Mode Network (DMN), PDMN = Posterior DMN, RFP = Right Fronto Parietal, LFP = Left Fronto Parietal, AUD = Auditory, TPJN = Temporo-Parietal Junction Network, LANG = Language Network, EXEC = Executive Control Network, INS = Insular Network, MSMN = Medial Sensory-Motor Network (SMN), VSMN = Ventral SMN, SSNR = Somatosensory Network - Right, SMNL = Somatosensory Network - Left, BGN = Basal Ganglia Network, SCRB = Superior Cerebellar Network, PCRB = Posterior Cerebellar Network; R = Right Hemisphere of the Brain, L = Left Hemisphere of the Brain.

Figure 2 A (top row) demonstrates regions with significantly enhanced FC in the COVID-19 group compared to the HC group for the *BGN* network. The FC of the *BGN* network was enhanced within the *Right – Calcarine Cortex (Calc), Cuneus (Cu*) and *Lingual Gyrus (LiG*) regions, comprising the *occipital* lobe. Similarly, the COVID survivors also demonstrated enhanced FC of the *PRN* network (Figure 2 B) with regions from the *Parietal Lobe: Bilateral – Superior Parietal Lobule (SPL*) and *Precuneus (PCu*) regions. On the other hand, for the *LANG* network, Figure 2 C shows reduced functional connectivity among COVID participants compared to HCs in several layers of the *cerebellar vermal lobules (CVL) (I-V, VI-VII*). The cluster peak information including peak *t*-scores and *FWE* corrected exact p-values with relevant anatomical regions from each network have been tabulated for an easy reference in *Table 3*.

**Figure 2.**
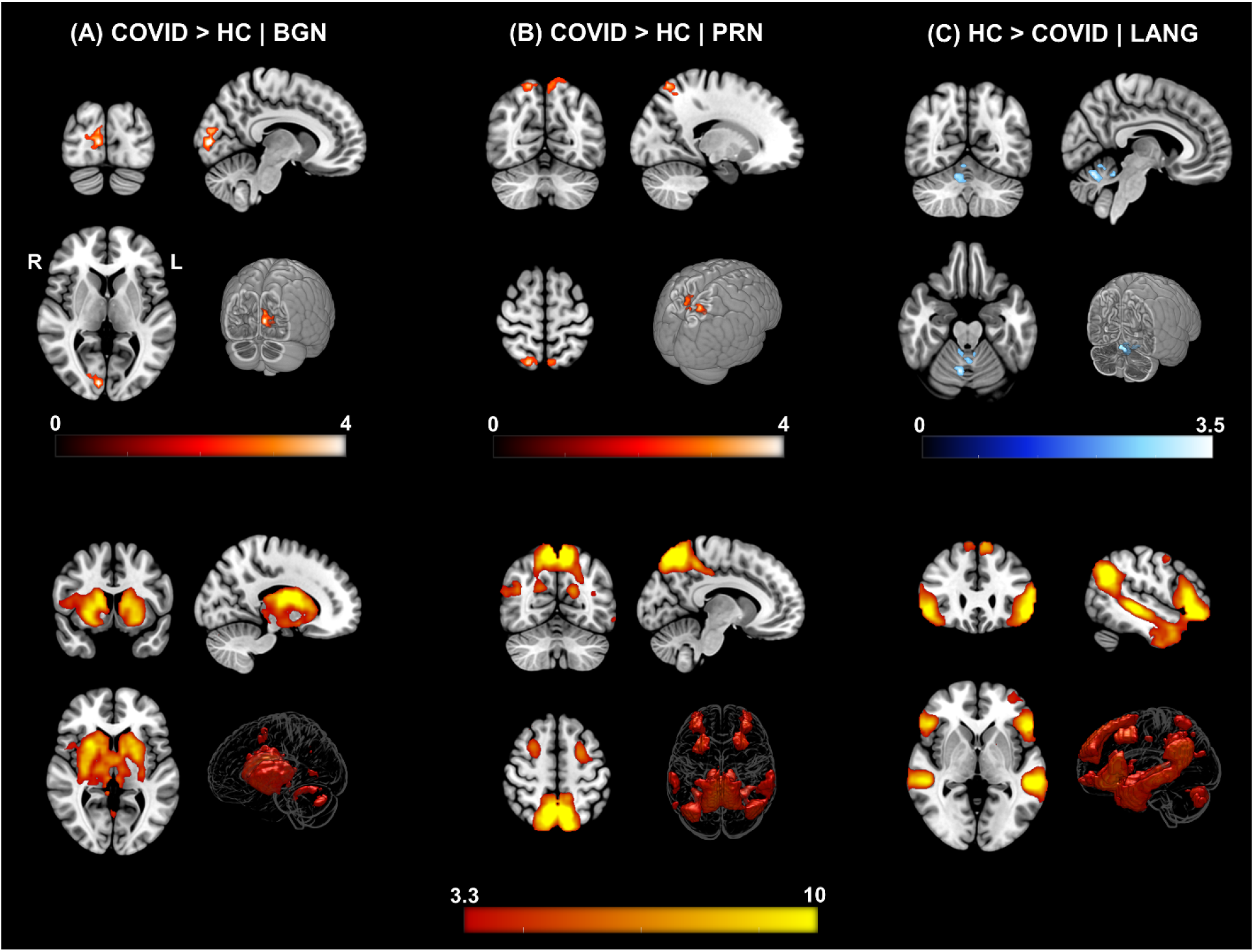
ΔFC | Functional Connectivity differences between COVID survivors and healthy controls. [top row] (A) COVID > HC: Enhanced FC in COVID compared to HCs observed in the *Basal Ganglia Network (BGN*) network.Three orthogonal slices (left) along with a cut-to-depth volume rendered image to show the effects in the *right Calc, Cu* and *LiG*. The colorbar represents *t*– score values. Cluster information include - cluster peak: *[9 −84 6]*, and cluster size *= 69 voxels*. **(B) COVID > HC:** Enhanced FC in COVID compared to HCs observed in the *Precuneus (PRC*) network, demonstrating a significant difference in FC in the *bilateral SPL and PCu* regions. Cluster information include - cluster peak: *[21 −57 54]* and cluster size *= 90 voxels*. Please note, enhanced FC among COVID survivors in both **(A)** and **(B)** is represented with a hot iron colormap and corresponding colorbar. **(C) HC > COVID:** Reduced FC in COVID compared to HC group observed in the *Language (LANG*) network demonstrating significant FC differences in several vermal layers of the *Cerebellum*. The electric blue colormap and corresponding colorbar are used to indicate that FC is reduced in the COVID group. Cluster information include - cluster peak: *[9 −63 −24]* and cluster size *= 57 voxels*. [**bottom row]** Corresponding group level ICA networks from which FC differences are shown on the top row. The colorbar represents z-scores.

**Table 3.**
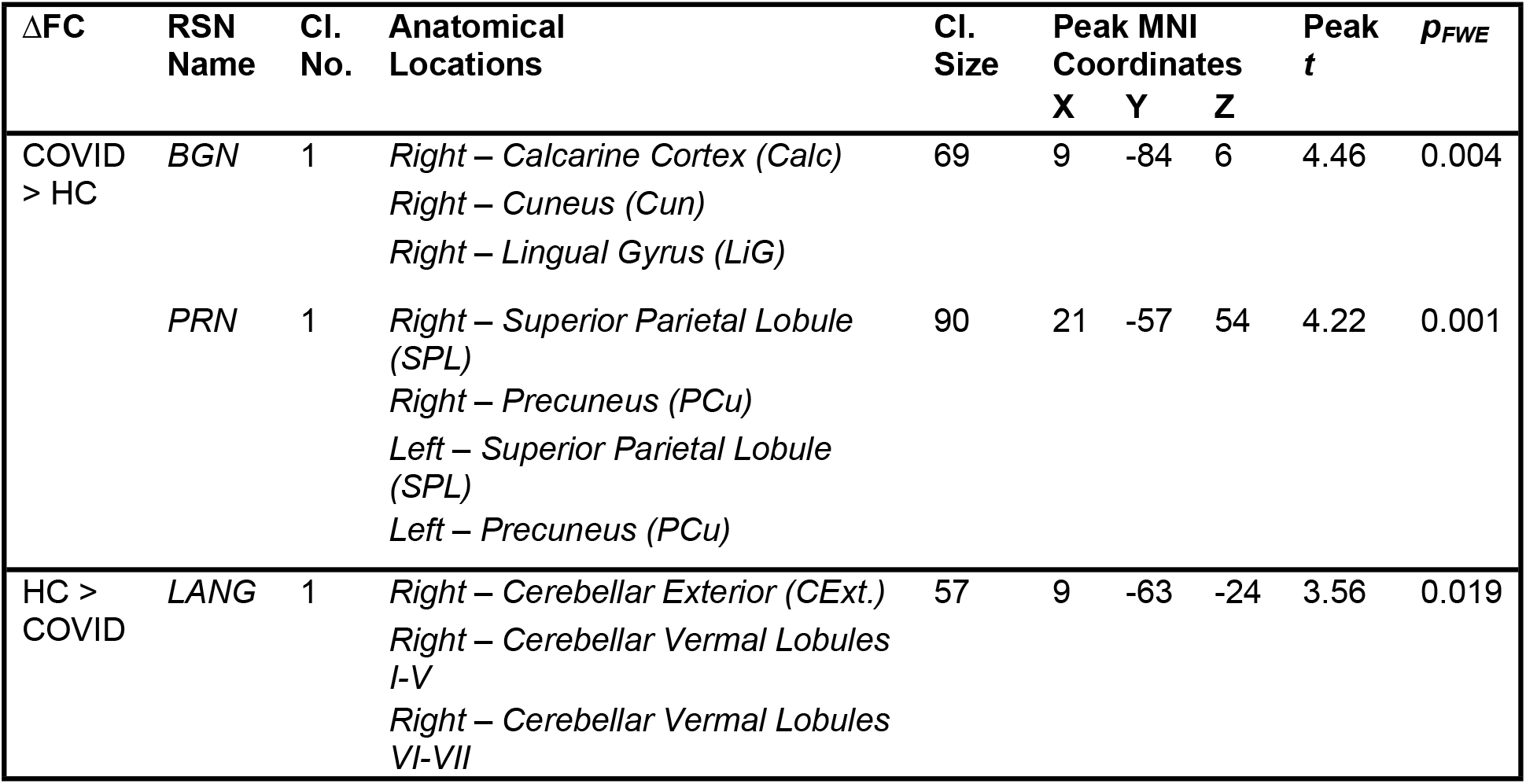
List of spatial regions from significant clusters obtained from the contrast – COVID > HC. The regions from three RSNs – *BGN, PRN* and *LANG* which demonstrated significant differences are presented with peak MNI coordinates (X Y Z) and corresponding *peak t*-score values for each cluster. **Keys:** ΔFC = Direction of change in Functional Connectivity; Cl. = Cluster; Cl. No. = Number of Clusters; Cl. Size = Cluster Size; Peak *t* =*peak t*-score; *p_FWE_* = family wise error corrected p-value.

| $\Delta FC$ | RSN Name | Cl. No. | Anatomical Locations | Cl. Size | Peak MNI Coordinates | | | Peak $t$ | $p_{FWE}$ |
| --- | --- | --- | --- | --- | --- | --- | --- | --- | --- |
|  |  |  |  |  | X | Y | Z |  |  |
| COVID > HC | BGN | 1 | Right – Calcarine Cortex (Calc)<br>Right – Cuneus (Cun)<br>Right – Lingual Gyrus (LiG) | 69 | 9 | -84 | 6 | 4.46 | 0.004 |
|  | PRN | 1 | Right – Superior Parietal Lobule (SPL)<br>Right – Precuneus (PCu)<br>Left – Superior Parietal Lobule (SPL)<br>Left – Precuneus (PCu) | 90 | 21 | -57 | 54 | 4.22 | 0.001 |
| HC > COVID | LANG | 1 | Right – Cerebellar Exterior (CExt.)<br>Right – Cerebellar Vermal Lobules I-V<br>Right – Cerebellar Vermal Lobules VI-VII | 57 | 9 | -63 | -24 | 3.56 | 0.019 |

Figure 3 shows brain regions where a significant negative correlation was observed between FC and self-reported fatigue, from the *PRN* network. The statistic map (Figure 3, *left*) shows the cluster where a negative correlation between FC of the PRC network and fatigue scores was observed in the *Left – Superior Parietal Lobule (SPL), Superior Occipital Gyrus (SOG), Angular Gyrus (AnG*) and *Precuneus (PCu*). The graph on the right visually portrays this negative relationship (*Spearman’s ρ = −0.47, p = 0.001, r^2^ = 0.22*) between the average FC of this cluster and fatigue scores. The scatter plot (Figure 3, *right*) clearly shows that the effects of FC and fatigue are significantly larger in the COVID group (light pink dots higher than cyan dots) compared to HC group. A more intuitive version of this figure can be found within the Supplementary Materials (see Figure S3) where the linear regression line within each group is demonstrated separately to clearly show that the COVID group was more susceptible to fatigue and majorly drove the overall linear trend shown in Figure 3. This can be observed from a significantly stronger correlation between FC and fatigue among survivors (*ρ = −0.72, p = 0.00002, r^2^ = 0.52*) compared to the HC group (*ρ = −0.36, p = 0.163, r^2^ = 0.13*).

**Figure 3.**
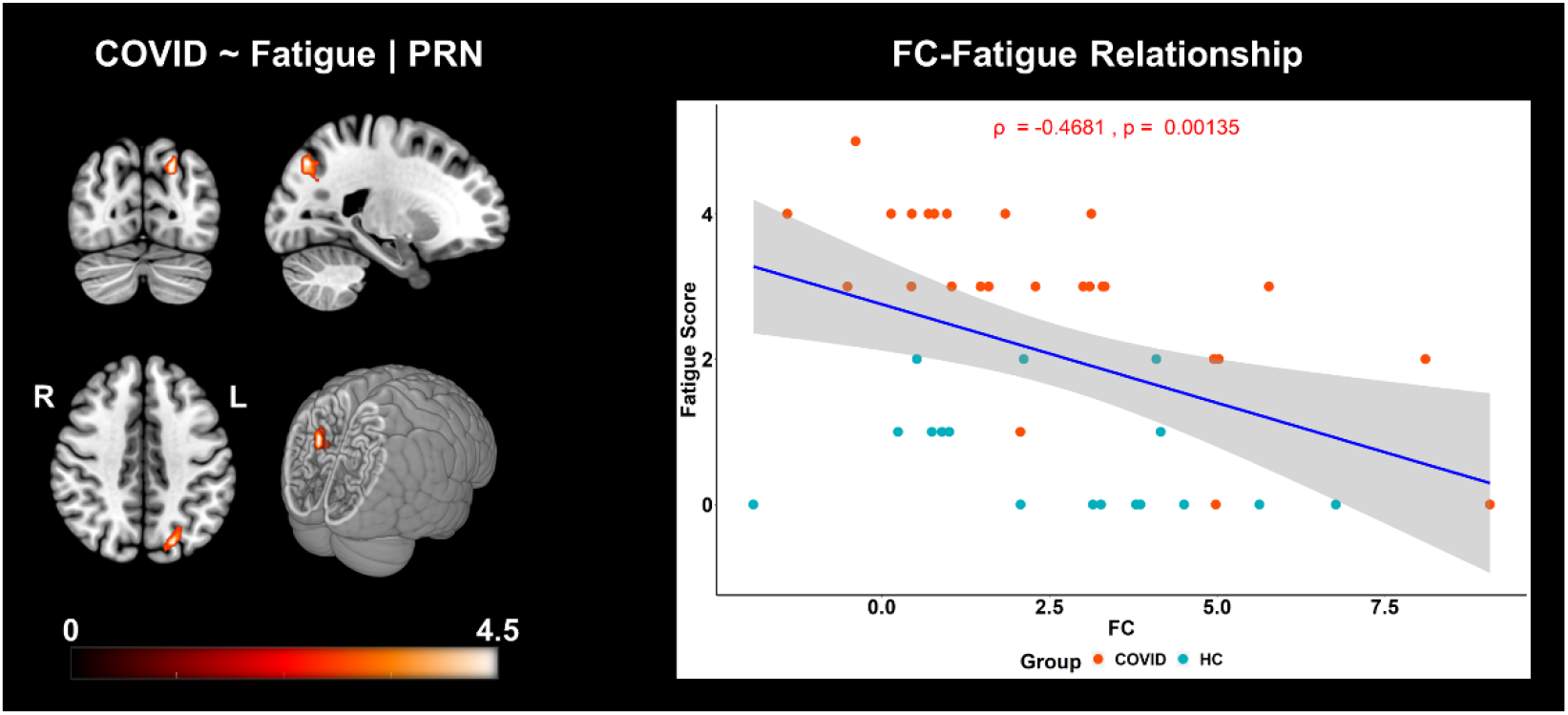
FC is negatively correlated with self-reported fatigue scores in COVID and HC individuals. The three orthogonal slices on the left, shows the cluster withing the *PRN* network, along with a cut-to-depth volume rendered image consisting of *Superior Parietal Lobule (SPL), Superior Occipital Gyrus (SOG), Angular Gyrus (AnG), Precuneus (PCu*). This cluster demonstrated significantly negative correlation with fatigue. The colorbar represents t-score values. The cluster consisted of *46* voxels and the peak was located at MNI coordinates:*[−21 −72 42]*. The peak *t*-score of the cluster was, *t_peak_ = 4.40*, and corrected for multiple comparisons by controlling false discovery rates, at *p_fdr_* < 0.05. The graph on the right shows the linear relationship between FC within the significant cluster and self-reported fatigue scores across all groups. The x-axis represents the residuals plus the average FC (z-scores) across groups from the cluster and the y-axis represents the fatigue scores. The light pink dots represent the COVID group, and the cyan dots represent the HC group. The shaded gray area represents the 95% confidence interval. The blue line represents the least squares regression line of best fit.

## Discussion

The results from this study support our hypothesis that COVID survivors demonstrate altered FC when compared to HCs, even two weeks after discharge from the hospital. Furthermore, the results from the work-related fatigue analysis support the hypothesis that the COVID survivors experience significantly higher fatigue during work and demonstrate more susceptibility to fatigue, with stronger negative correlation of FC and fatigue compared to HCs. Our hypothesis on FC alterations was based on both early case-reports and more recent group level neuroimaging reports of structural and functional brain alterations. Individual case reports were primarily from acutely ill patients using FLAIR (Kandemirli et al., 2020; Kremer, Lersy, Anheim, et al., 2020; Paterson et al., 2020) and Susceptibility Weighted Imaging (SWI) (Conklin et al., 2021), whereas, group level reports, such as those derived from fMRI, include, reduced *default mode* and *salience* connectivity (Fischer et al., 2021) and high prevalence of abnormal time varying and topological organizations between *sensorimotor* and *visual* networks (Fu et al., 2021).

In the current context, we report between group FC alterations of three large scale RSNs –*BGN*, *PRN* and *LANG* networks and further show stronger negative correlation between FC in the *PRN* network with self-reported fatigue at work in COVID survivors compared to the HC group.

We observed enhanced FC of the *BGN* network in the COVID survivors compared to the HCs within *Calc, Cu* and *LiG. Calc* and *Cu* are primarily involved in visual processing. Fu and colleagues reported that COVID survivors demonstrated enhanced connectivity between *Cerebellum, Sensorimotor* and *Visual* networks, characterizing that they spent abnormally higher time in a specific brain state compared to healthy controls (Fu et al., 2021). A recent study has also suggested *Cu* to be a major hub for mild cognitive impairment in idiopathic REM sleep behavior disorder (iRBD) (Mattioli et al., 2021). *LiG* and weak *insular* coactivation with the *occipital* cortex have been shown to be associated with disrupted salience processing that can lead to loss in motivation in day-to-day tasks (Kim et al., 2018). Interestingly, cortical thickness alterations were also reported in *Calc* and *LiG* regions in non-hospitalized and mildly symptomatic survivors (Crunfli et al., 2021). In our cohort of hospitalized survivors, we expect these alterations to scale up with severity, as severity tends to increase chances of neurological manifestations in hospitalized COVID-19 survivors (Mao et al., 2020). Moreover, the *basal ganglia* are known to be associated with fatigue (Miller et al., 2014), cognitive, emotional and attention processing (Di Martino et al., 2008; van Schouwenburg et al., 2015). The synergy of these studies to our findings indicates possible functional brain associations of commonly observed symptoms in survivors with post-acute sequelae SARS-CoV-2 infection (PASC or Long COVID) lasting many months (Carfì et al., 2020; Garrigues et al., 2020; Logue et al., 2021; Peluso et al., 2021).

We observed enhanced FC of the *PRC* network among COVID survivors compared to the HC group. Enhanced FC in this network was observed in the *Bilateral SPL* and *PCu* regions*. PCu* is a constituent of *DMN*, and higher functional connectivity with this region may indicate some compensatory mechanism due to loss in connections in other pathways. Moreover, alterations between *DMN* and *salience* connectivity has been recently reported in a follow-up study from Fischer and colleagues (Fischer et al., 2020), although, initially, a single patient showed no differences in FC of *DMN* when compared to five healthy controls (Fischer et al., 2020). *PRN* network consists of *Precuneus (PCu), Frontal Eye Fields (FEF*) and parts of the *Superior Parietal Lobule (SPL). SPL* is a constituent of the *posterior parietal cortex (PPC*) which has been shown to have functional association with altered *anterior insula* connectivity in CFS (Wortinger et al., 2017). Moreover, these brain regions are also known to be involved in attention processing, therefore, enhanced FC in these regions may indicate possible compensatory mechanisms of attention related symptoms that recovering patients may experience. This is significant, because about 37% of the COVID survivors in our study reported lack of attention and all these 37% of the participants also reported a work-related fatigue score of 2 or higher on a scale of 5. Therefore, further investigations are necessary to understand these processes better, especially, from a clinical perspective.

We also observed reduced FC of the *LANG* network within several layers of the *Cerebellar Vermal Lobules* among COVID participants when compared to HCs. These lobules have been suggested to be involved in cognition and emotion processing (Park et al., 2018). Interestingly, structural alterations have also been reported in these layers among COVID survivors. A case report of a 47-year-old male described hyperintense *bilateral cerebellar hemisphere* and *cerebellar vermis*, which was also the first reported case of acute cerebellitis in COVID-19 (Fadakar et al., 2020). Another case of cerebellitis was also reported recently, adding on to the wide range of neurological disturbances in the CNS (Malayala et al., 2021). Moreover, approximately 45% of the survivors in the study experienced loss/reduction of sense of smell or hyposmia. Activation in the *cerebellum*, specifically in the *vermal lobules*, through olfactory stimulation has been shown both in humans (Ferdon & Murphy, 2003) and animals (García et al., 2015). Reduced FC within *cerebellar vermal layers* may indicate connectivity deficits, as a result of olfactory dysfunction.

We observed FC alterations in three out of five networks of interest. The neuro-invasion of the coronavirus can spread and sustain throughout the brain, causing pathologies several months after the initial infection (Daniel et al., 2022), possibly leading to development of PASC. (Daniel et al., 2022) also found traces of the virus in brain tissue biopsies even from mild and asymptomatic patients. Therefore, perhaps with time, more and more structural and functional networks in these survivors will be affected during the transition phase to PASC. With the availability of more sample data and a follow-up design, perhaps all brain networks can be investigated for a more comprehensive understanding of the brain pathologies systematically. RS-fMRI can be quite useful in this regard because it facilitates the investigation of multiple brain networks across the whole brain (Damoiseaux et al., 2006; Raichle & Mintun, 2006; Shulman et al., 2004). The results also suggest a possible link between structural and functional abnormalities in COVID survivors since the FC alterations were observed in regions that align with anatomical regions exhibiting hyperintensities, particularly, in the *basal ganglia* (Paterson et al., 2020), *parietal* and *occipital* lobes (Kandemirli et al., 2020) as well as in *cerebellar* regions (Kremer, Lersy, de Sèze, et al., 2020). We do not know how the functional abnormalities in these survivors relate to PASC, however, cerebrovascular injuries and inflammatory processes may play an important role in determining whether a patient returns to normal health or continues ill with PASC. Neurological damage and abnormalities found in cerebral arteries in several acute patients across various centers (Gulko et al., 2020), can imply different levels of cerebral blood flow alterations in PASC and not-PASC survivors. Thus, it is possible that functional changes will be observed between PASC and not-PASC COVID survivors in future studies.

Based on recent literature on PASC patients, fatigue has been the most frequently reported symptom (Logue et al., 2021; Peluso et al., 2021; Tabacof et al., 2020). In the current study, we also observed that fatigue during work was the highest reported symptom (86.84%) among COVID survivors 2 weeks after hospital discharge. We observed higher fatigue levels among COVID survivors when compared to healthy controls (*p < 0.001*). We further evaluated linear relationship between FC of the *PRC* network and self-reported fatigue at work across both COVID and HC participants. We observed a significant negative correlation of FC with fatigue within the *Left SPL, SOG, AnG* and *PCu*, i.e., brain regions primarily belonging to the *parietal* lobe (see *Table 4* for cluster information). Interestingly, our recent investigation using the same group of survivors revealed stronger positive correlation between GMV and self-reported fatigue within the *precuneus* and *SPL* regions, when compared to HCs (Hafiz et al., 2022). This provides supporting evidence to our hypothesis that COVID survivors also demonstrate stronger correlation between FC and fatigue in the *PRC* network, based on our previous observation on GMV and fatigue relationship in these regions. Moreover, structural atrophy in the *parietal lobe* has been shown to be associated with fatigue among multiple sclerosis (MS) patients (Calabrese et al., 2010; Pellicano et al., 2010). An RS-fMRI study involving CFS patients, used ICA to reveal loss of intrinsic connectivity in the *parietal lobe* (Gay et al., 2016). Therefore, the fact that lower FC in the *parietal lobe* correlates more negatively to higher fatigue scores among COVID survivors, can be clinically relevant because it matches both structural and functional relationships with fatigue in other neurological disorders such as MS and CFS. To the best of our knowledge, this is the first study to show work-related fatigue correlates of FC among recovering patients 2 weeks after hospital discharge. Therefore, future studies are necessary to evaluate this highly prevalent symptom further in the surviving cohorts.

**Table 4.**
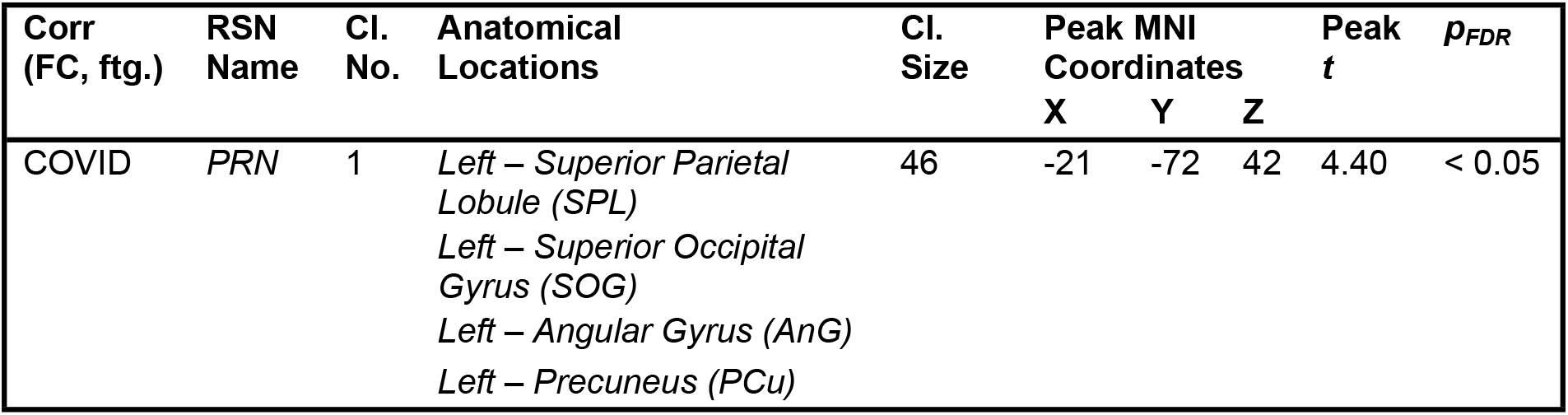
List of spatial regions from the cluster showing significant correlation with selfreported fatigue among COVID individuals. The regions from *PRN* which demonstrated significant correlation are presented with peak MNI coordinates (X Y Z) and corresponding peak *t*-score values for each cluster. **Keys:** FC = Functional Connectivity; ftg. = Fatigue Scores, Cl. = Cluster; Cl. No. = Number of Clusters; Cl. Size = Cluster Size; *t* = peak *t*-score; *p_FDR_* = false discovery rate corrected p-value.

## Limitations

Despite our efforts to show group level effects that reflect individual and group level reports in the recent literature, our study still maintains a cross-sectional design. In cases like this, a better approach for the future would be to use follow up designs (Fu et al., 2021; Lu et al., 2020; Tu et al., 2021) or possibly a longitudinal design where patients could be observed both before and after the pandemic like the one using the UK-biobank (Douaud et al., 2021). Our effort here, was to show group level effects at an early stage of recovery (2 weeks after hospital discharge) and determine the relation between work-related fatigue and FC of RSNs. We believe the results from this study will help understanding the recovery stage brain alterations and how they might drive fatigue-related symptoms among COVID survivors.

## Author Credit Statement

**Rakibul Hafiz**: Methodology, Software, Formal Analysis, Data Curation, Writing – Original Draft, Review and Editing.

**Tapan K. Gandhi:** Conceptualization, Investigation, Resources, Supervision, Writing – Review and Editing.

**Sapna Mishra:** Investigation, Resources, Data Curation, Writing – Review and Editing.

**Alok Prasad:** Writing – Review and Editing.

**Vidur Mahajan:** Writing – Review and Editing.

**Xin Di:** Methodology, Writing – Review and Editing.

**Benjamin H. Natelson:** Writing – Review and Editing.

**Bharat Biswal:** Conceptualization, Resources, Project Administration, Supervision, Writing – Review and Editing.

## Acknowledgements

This study was supported by NIH grants: R01AT009829 and R01MH131335 and MeitY (Government of India) under Grant No: 4(16)/2019-ITEA

