## Supplementary Materials for "Assessing functional connectivity differences and work-related fatigue in surviving COVID-negative patients"

**Figure S1.** **Participant recruitment questionnaire for symptoms during hospitalization.** The figure shows the questions that the participants were asked to answer to evaluate the development and duration of specific symptoms relating to COVID-19 during hospitalization.

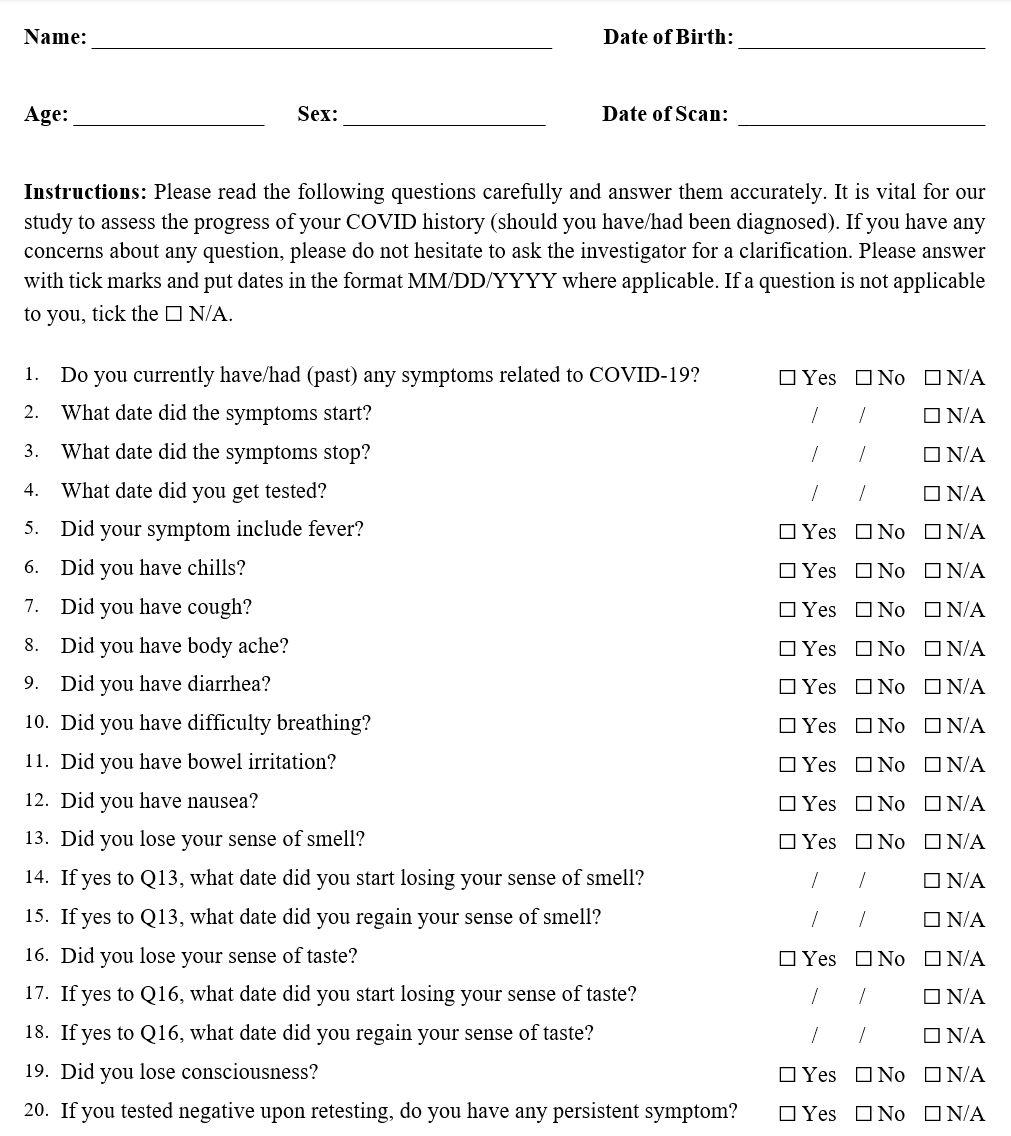

**Figure S2.** **Fatigue and long-COVID related symptoms questionnaire.** The ‘Part 1’ on the left is the fatigue questionnaire based on ‘Life Spheres Criteria’ shown in the bullet points. The participants were asked to grade their fatigue levels on a scale of 0-5 with increasing fatigue severity as the number approaches 5. A secondary, ‘Part 2: Symptoms Criteria’ questionnaire (on the right) was also provided to the surviving patients. This was done to identify persistent or new symptoms possibly relating to PASC development. We were particularly focused on the fatigue levels experienced during work and all results and evaluations are reported based on the fatigue scores during work.

Source: Natelson, B. H. (2019). Myalgic Encephalomyelitis/Chronic Fatigue Syndrome and Fibromyalgia: Definitions, Similarities, and Differences. Clinical Therapeutics, 41(4), 612-618. https://doi.org/https://doi.org/10.1016/j.clinthera.2018.12.016

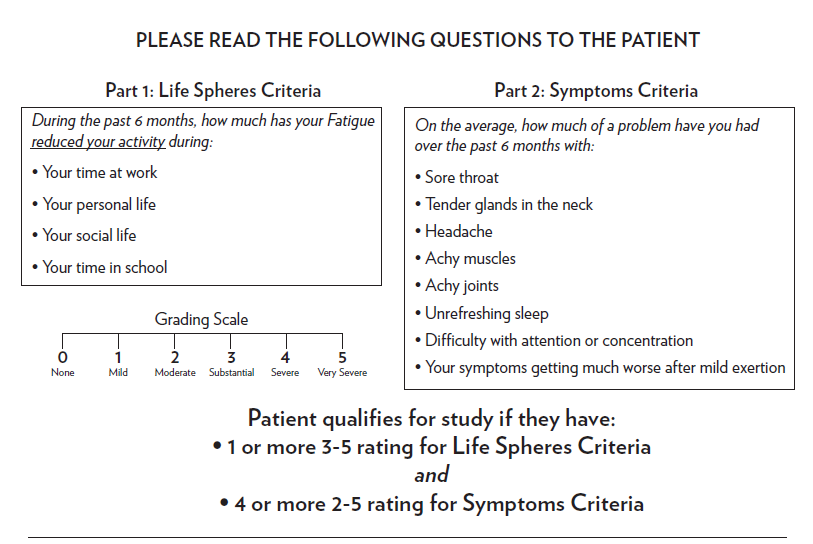

**Figure S3.** Similar to Figure 3 in the revised manuscript, except, here we show the correlation and linear regression line for each group in a single plot to better demonstrate the difference in effect for the significant cluster. The orange dots and regression line representing the COVID group clearly demonstrate stronger effects of FC and fatigue compared to HC group.

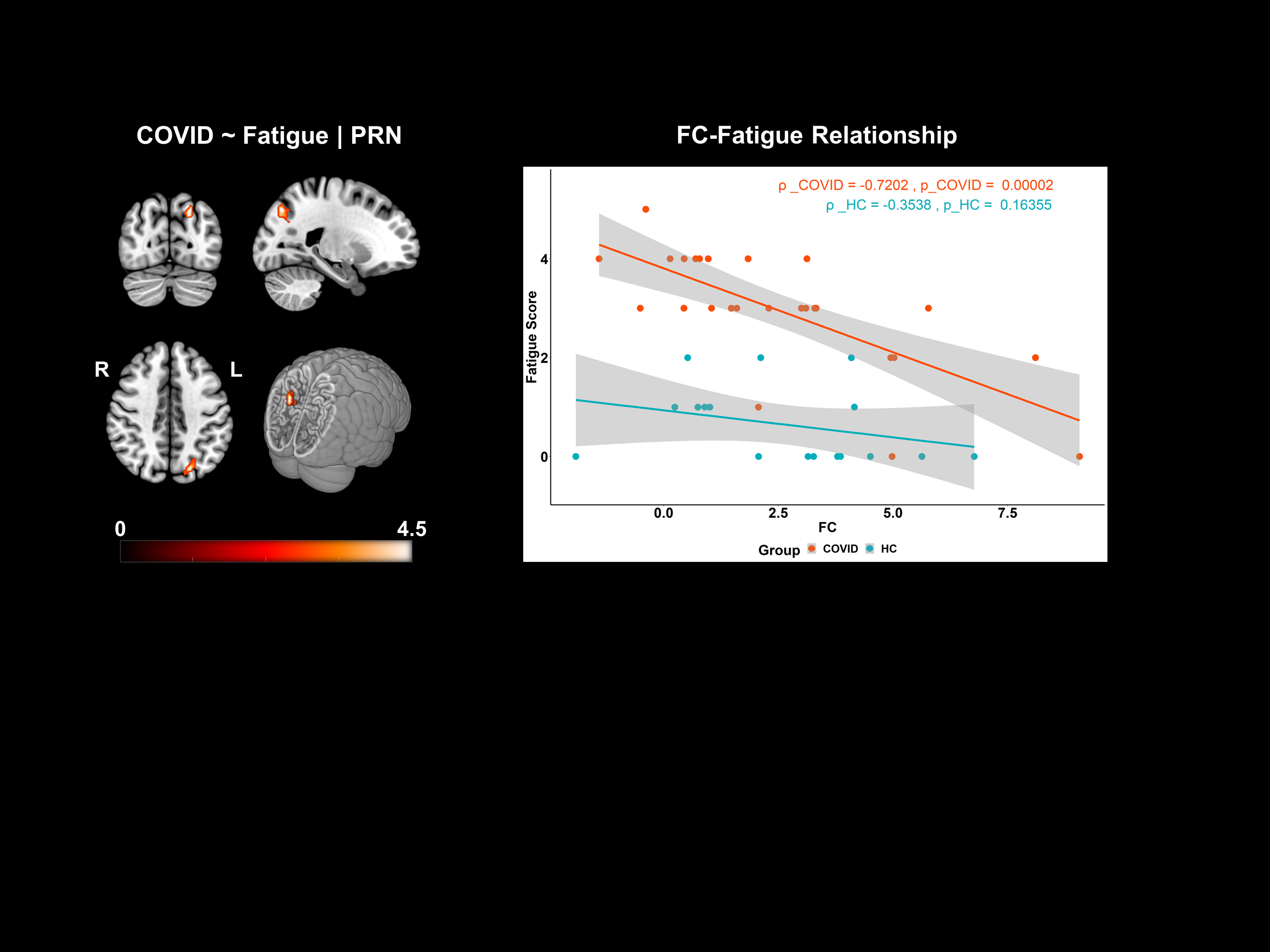
